# Language-Driven Clustering of Transfer, Persistence, Prevalence & Recovery Studies for Forensic Activity-Level Evaluation

**DOI:** 10.64898/2026.11.11.766034

**Authors:** Janet Stacey, Stephanie Opperman, James M. Curran, Sally Ann Harbison

## Abstract

Transfer, Persistence, Prevalence, and Recovery (TPPR) data are essential for forensic activity level evaluations (ALE); however, concerns remain regarding the accessibility of relevant literature. This study investigated whether automated literature retrieval and natural language processing (NLP) methods could be used to identify and characterize TPPR studies. Existing forensic repositories were combined with records retrieved through the Web of Science Application Programming Interface (API) and enriched with bibliographic metadata, producing a corpus of 3,279 unique records. Automated retrieval tools were evaluated, and transformer-based topic modeling was applied to examine the breadth of the literature. The results showed that database APIs provided the most reliable mechanism for literature retrieval, while topic modeling identified meaningful groupings corresponding to major forensic disciplines and trace evidence types. The findings indicate that TPPR literature exists and that automated methods can support its discovery and organization. Although the availability of literature does not necessarily imply suitability for ALE, the results suggest that the principal challenge is the discovery, assessment, and reuse of published studies rather than their absence. Automated retrieval, semantic search, and quality assessment frameworks may improve access to relevant evidence and support the development of forensic knowledge systems for evaluative reporting and Bayesian network construction.

## Introduction

Concerns have been raised in the ongoing discussion of the need for Activity Level Evaluations (ALE) regarding the lack of suitable data (Onorato, 2025; Hunt, 2025; Stacey et al., 2025). A requirement for conducting ALE is that it is underpinned by data relevant to the questioned activity. Determining appropriate search terms and trawling through public, indexed databases for supporting data can be time consuming. This cost may result in practitioners relying on previously found papers and being unaware of new pertinent information. A more systematic approach has been recommended for the recovery and assessment of Transfer, Persistence, Prevalence, and Recovery (TPPR) data (Gosch and Courts, 2019). Implementing such an approach may be difficult and time consuming to achieve. The inconsistency in the knowledge of different experts also makes it difficult to obtain a consensus among the forensic community. This may lead to disagreements between experts in the courtroom, thus complicating the decisions to be made by the fact-finder.

Information about published studies has been collected to simplify access to literature, create a standardized awareness of what is available, and support systematic searches. Some individuals and organizations have collected private file repositories of published TPPR studies for use in their evaluations. Gosch and Courts (2019) created a publicly available spreadsheet database resource called the *DNA-Trace Activity Database* (DNA-TrAc) for forensic biology cases. It is a collection of summary information about published DNA TPPR studies that allows practitioners to familiarize themselves with the existing literature on the topic. However, contributing to DNA-TrAc is voluntary, so the collection is limited, with updates occurring sporadically. The Forensic Science Research Group (GRSF) at the University of Quebec in Trois-Rivières, Canada, has also developed a TPPR knowledge base—called the *Transfer Traces at Activity Level Database* (TTADB)—for several trace types (Cadola et al., 2021). This database is maintained through student projects and the voluntary submission of articles by the forensic community. Both of these attempts to collate TPPR data in forensic-specific repositories, unfortunately, rely on manual search mechanisms to find relevant papers and are time consuming to maintain, with a risk of being abandoned. A comprehensive assessment of the current repositories of published TPPR studies is needed to determine what information has been gathered, how complete it is, and whether these resources are suitable for ALE.

Other criticisms of the current TPPR literature raised by members of the forensic and legal communities are that inter-laboratory variation is often too large to allow the use of results from other laboratories, and that the current literature often does not provide enough information to standardize TPPR findings for ALE use (Gill et al., 2025). Gill et al. (2026) published a set of minimum requirements for forensic genetic studies. The authors believe that if these requirements are met, then this will ensure that a study’s TPPR findings can be used by other laboratories.

Computer-aided retrieval of relevant TPPR data is an essential but currently missing component of fast and robust ALE. TPPR data repositories need to be kept up to date without manual curation; therefore, automated searching and retrieval of the literature are required. In recent years, there has been a growing number of tools that offer an automated literature search feature. Hewarathna A. I. (2023) has provided code that relies on open-source software tools to obtain data from various journal databases. Specifically, they provide use cases for IEEE Xplore, ScienceDirect, and ACM Digital Library. Clarivate, the owner of the Web of Science Online Journal Database, offers an Application Programming Interface (API) for programmatic access to search functionality (Clarivate, 2025). Lu et al. (2024) described an AI model that could perform the entire research process. The authors called this an *AI scientist*. The model is able to carry out literature searches in the field of computer science, iteratively improve current research, and write the experiment as a journal article for publication submission. This was followed in 2025 by the publication of Kosmos, an AI scientist algorithm designed for drug discovery (Mitchener et al., 2025). Additionally, a new generation of search tools, coined AI Research Assistants, has arisen. Some of the more commonly used tools of this type are Elicit (Elicit, 2023), Semantic Scholar (The Allen Institute for Artificial Intelligence, 2023), and Perplexity (Perplexity AI, 2026). Elicit and Semantic Scholar both utilize semantic similarity to search through curated datasets of millions of published articles. Semantic similarity is a metric that measures how closely two pieces of text, words, or concepts match in meaning rather than in exact wording (Miner et al., 2012). Perplexity uses large language models to collect data directly from websites across the internet. This latter method has been controversial, with copyright infringement lawsuits recently plaguing Perplexity AI (BakerHostetler, 2026; Leppard, 2025; Palmer, 2026; Metz and Grynbaum, 2025). Another potential hurdle is that forensic publications are most often published in subscription journals with paywalls. This makes them less accessible to these tools. A less pressing but still important concern is that these search tools have not been specifically assessed with the needs of forensic science in mind.

In this paper, the viability of these automated literature search tools is investigated for use in a forensic context. The contents of existing forensic datasets were also investigated, and the advantages and disadvantages of the different repositories and the data held by each are described. A natural language processing (NLP) model was developed, optimized, and tested to analyze trends and characteristics of the studies from the various data sources to describe the depth and breadth of TPPR data available for ALE. NLP is the use of computational methods to analyze items based on the terms, structure, and context of the language used. It looks for common words or other elements to find relationships between items. When those relationships are used to characterize them into clusters or topics, the technique is known as Topic modeling (Miner et al., 2012).

## Methodology

This study was divided into three main phases: First, a selection of available automated literature search tools was evaluated to determine their suitability for identifying TPPR studies relevant to forensic activity level evaluations. Second, multiple existing TPPR literature repositories were identified, accessed, and combined into a unified dataset with data retrieved by the search tools. Finally, the resulting corpus was cleaned, standardized, enriched with additional bibliographic metadata, and analyzed using natural language processing (NLP) techniques to investigate the breadth and characteristics of the available literature.

### Evaluation of automated literature search tools for a forensic use-case

Four types of automated literature search tools were investigated: (i) LiteratureReview: an open-source tool based on Selenium, (ii) the AI Scientist framework and its use of Semantic Scholar, (iii) an API to access the Clarivate ScienceDirect database, and (iv) direct web access to Semantic Scholar.

#### The open-source LiteratureReview tool

LiteratureReview (Hewarathna A. I., 2023) is an open-source software tool written in Python that collects data from various supported science journal databases. Supported databases are currently limited to IEEE Xplore, ScienceDirect, and the ACM Digital Library. The tool relies on other open-source python software tools to navigate to, search, and return the web content, i.e. HTML, from a search results webpage. The code for the tool is available on GitHub. The tool was downloaded from the code repository into a Python 3.13.5 environment for testing.

#### The AI Scientist model and Semantic Scholar

The AI Scientist (Lu et al., 2024) is another open-source, Python-based software tool. The tool itself relies on the Semantic Scholar API (The Allen Institute for Artificial Intelligence, 2023), access to a computer running a Linux-based operating system equipped with a graphics processing unit (GPU), and access to a Large Language Model (LLM)—either locally or via API access to a commercial provider such as OpenAI or Anthropic. The tool covers the complete scientific experimentation process, from literature review to writing publications.

#### The Clarivate API

Clarivate offers two APIs for the Web of Science indexed database: the Web of Science Starter API and the Web of Science API -Expanded. The use of an API allows automated access to search results. As the Web of Science API -Expanded option is a paid subscription, the Starter API was tested in this study. This API was accessed using a Python v3.13.5 script and a trial API key provided by Clarivate. The query string that was used is shown in Figure 1. This query contained several conditions to focus results on TPPR studies by filtering out terms related to Digital Forensics, Forensic Psychology, and Astrophysics. Other optional settings that were used for the query are shown in Table 1. The fields returned from the query are listed in A.

**Figure 1:**
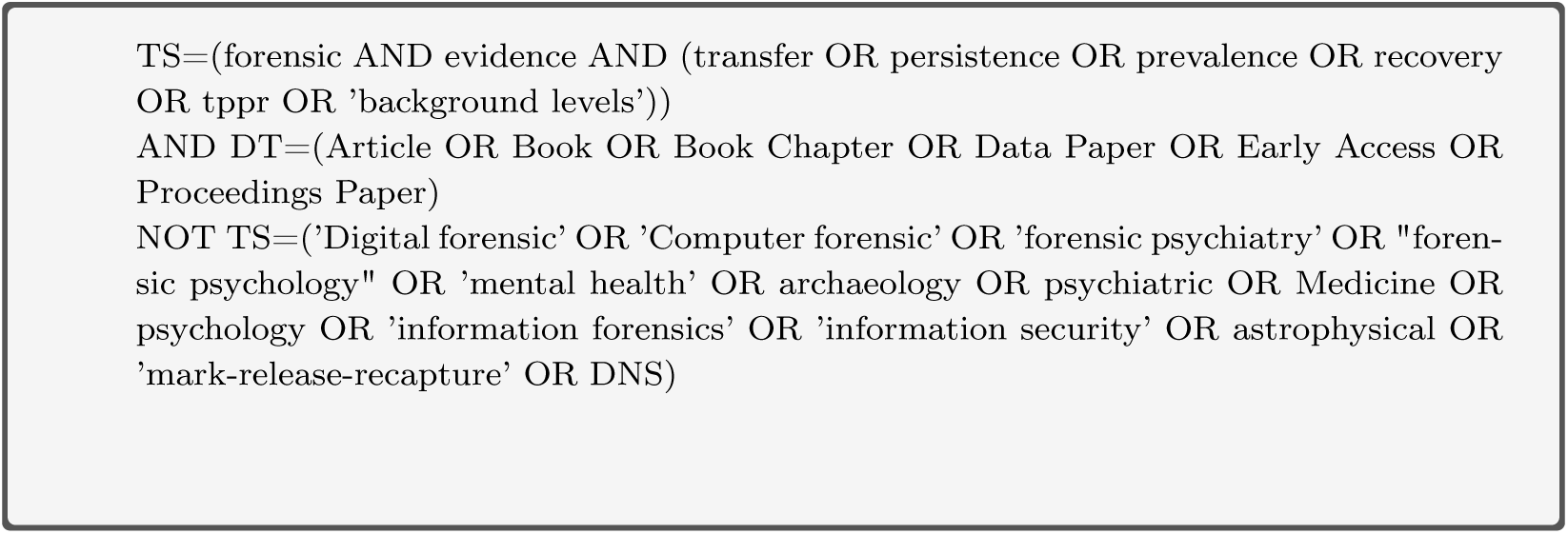
Character string used for Web of Science API Post Request. Field names are as follows: TS=Topic, DT=Document Type

**Table 1.** Settings used for Web of Science Starter API Post Request

| **Argument Name** | **Details** | **Value** |
|---|---|---|
| db | Web of Science Core collection | WOS |
| limit | set the limit of records on the page (1-50) | 50 |
| page | set the number of returned pages (20) | range(1,21) |
| sort_field | Order by field(s). LD for Load Date, D for Descending order | LD+D |
| modified_time_span | Defines a date range in which the results were most recently modified | None |
| tc_modified_time_span | Defines a date range in which times cited counts were modified | None |
| detail | Returns the full data by default | None |

The responses received from the API request were combined with data from the existing forensic repositories for analysis.

#### Direct Access to Semantic Scholar

Semantic Scholar (The Allen Institute for Artificial Intelligence, 2023) was also accessed directly through the web interface rather than via an API. It was initially provided with the same search query as the Web of Science API (Figure 1), and then reformatted to fit the requirements of the tool. The results that were returned were assessed to determine their suitability for inclusion.

### TPPR data sources

Five different sources of bibliographic information relating to TPPR studies were identified for inclusion in the analysis. These sources were included because they are commonly used and publicly accessible. Although each repository was developed independently, it was anticipated that they would contain overlapping bibliographic and descriptive information. Combining information from multiple sources was, therefore, expected to provide a richer and more complete description of each published study.

These sources differed in both their origin and storage format, comprising collections of file-based PDF articles, an existing BibTeX bibliography, and two curated databases. To enable consistent downstream processing, bibliographic information from all sources was converted to a common format prior to analysis.

These sources are:

*•* file collection shared from the Faculty of Law, Criminal Justice and Public Administration at the University of Lausanne (77 articles, 11 theses)
*•* TPPR papers discovered during the literature review (58 articles) (Stacey et al., 2025)
*•* John Butler forensic biology bibliography (751 records) (Butler and Willis, 2020; Butler, 2023, 2026).
*•* DNA-TrAc database (311 records) (Gosch and Courts, 2019)
*•* Transfer Traces at Activity Level Database—TTADB (2438 records) (Cadola et al., 2021)

### Data Preparation

The data from each of the data sources was read into a Python v3.13.5 environment for analysis. Each dataset was cleaned to ensure consistency in data type, column names, and content format. Empty and/or irrelevant columns were removed before the data was merged into a single dataframe. Specifically, the data cleaning carried out in Python and Microsoft Excel consisted of:

1. changing the data encoding to Unicode Transformation Format — 8-bit (UTF-8) (Yergeau, 2003), a commonly-used character encoding system that is tolerant of characters specific to different languages, such as those in non-Latin languages, diacritical marks, and other symbols. Some of these characters were not received correctly and required manual correction;
2. extracting and separating bibliographic information into individual metadata fields, including author, title, publication year, journal, conference, and publisher;
3. standardizing bibliographic fields to remove typographical inconsistencies and ensure consistent formatting;
4. identifying duplicate publications and retaining the most complete record;
5. converting all metadata fields to character format and normalizing text to facilitate dataset integration; and
6. merging the cleaned datasets into a single bibliographic database.

Many of the records did not include a unique persistent identifier, such as a Digital Object Identifier (DOI). This required data to be merged using the Article Title, Authors, Publication Title, and Year of Publication fields instead. An API from crossref.org (Hendricks et al., 2020) was later used to enrich the records’ metadata, allowing the association of a DOI to each article by title .

All data were manually curated to ensure accuracy and to remove duplication. Columns containing information relevant to the article content were concatenated into a single corpus of text for each record and analyzed using topic modeling.

Topic modeling (Blei et al., 2003) was carried out to determine commonalities in content characteristics using BERTopic v0.17.4 (Grootendorst, Maarten, 2025). The analysis methodology used followed that of Weng, Min-Hsien et al. (2022). The components used for analysis were embedding, dimensionality reduction, clustering, topic determination, and topic representation.

#### Embedding

Embedding is a method used to assess records for their semantic textual similarity to each other. It does this by encoding them as numbers and then using a similarity metric, usually cosine similarity (Tan et al., 2019), to collate them into topics. A transformer-based embedding model was used, specifically a Python implementation (sentence-transformers v5.5.1) (Aarsen, Tom, 2025) of a pretrained Sentence Transformer model called SBERT (Reimers, Nils and Gurevych, Iryna, 2019). Two base models were trialed: the default *all-mpnet-base-v2* model and the *paraphrase-multilingual-mpnet-base-v2* model. The latter was used as it is a small model with good general performance for multiple language inputs (Ubiquitous Knowledge Processing (UKP) Lab, 2025). Cosine similarity was used as the similarity metric. The following prompt was used: “Identify the topic or theme of forensic sample type or discipline based on the provided text. Do not include law enforcement, justice, statistics or type of crime as themes. Label topic with trace type or originating location”.

#### Dimensionality Reduction

Dimensionality reduction is the process of reducing the number of variables (features) used to represent a dataset while retaining as much of the important information as possible. It is a common preprocessing step in natural language processing because each unique word in a corpus is typically represented as a separate feature in the document-term matrix. As the vocabulary size increases, the resulting feature space becomes increasingly sparse, making it more difficult to identify meaningful similarities between documents and increasing computational complexity. This phenomenon is commonly referred to as the *curse of dimensionality*.

Uniform Manifold Approximation and Projection (UMAP) was used to project the high-dimensional document vectors into a lower-dimensional embedding while preserving the local neighborhood structure of the data. The size of the neighborhoods (n_neighbors) of 15 and 20, the embedding dimensions (n_components) of 2 and 5, and the use of cosine distance as a similarity metric were evaluated during model optimization (Section 2.4). Because UMAP is a stochastic algorithm, the random seed (random_state) was fixed at 38 to ensure reproducibility.

#### Clustering

Clustering is an unsupervised learning technique that groups documents, in this case, TPPR references, into clusters where documents within a cluster are more similar to each other than to documents in other clusters. Two complementary approaches to performing clustering were investigated: Zero-Shot Topic Modeling and Hierarchical Density-Based Spatial Clustering of Applications with Noise (HDBSCAN).

Zero-Shot Topic Modeling assigns documents to a predefined set of candidate topics where sufficient semantic (meaning-based) similarity exists. Documents that cannot be confidently assigned to one of these predefined topics are subsequently grouped using unsupervised clustering based on semantic similarity. BERTopic was provided with a list of 26 candidate topics, as shown in C. A minimum similarity threshold for inclusion was set at 0.8 (zeroshot_min_similarity parameter).

Hierarchical Density-Based Spatial Clustering of Applications with Noise (HDBSCAN) is an unsupervised, density-based clustering algorithm that automatically identifies clusters of varying sizes and shapes without requiring the number of clusters to be specified *a priori*, making it well suited for exploratory topic discovery (Campello et al., 2015). The Python implementation of HDBSCAN (v0.8.44) from the Scikit-Learn ecosystem (v1.9.0) was used (Campello et al., 2013). HDBSCAN was selected for this study because the number of topics present in the corpus was unknown.

#### Topic Determination

Topics are described by their top occurring terms. Determination of these is carried out during the embedding process, specifically by vectorization. Vectorization is the process of converting text into a numerical representation that a machine learning or NLP algorithm can work with. By default, this is performed using Term Frequency-Inverse Document Frequency (TF-IDF). TF-IDF converts the text into a sparse matrix of terms based on their observed frequency in the records. The most frequent terms in the documents are used to describe the topics. BERTopic uses the CountVectorizer function from the Scikit-Learn module (v1.9.0) to perform TF-IDF. The parameters used for CountVectorizer were changed to a term size of 1-4 ngrams and a term document frequency of between 0.2 and 0.6. These parameters were chosen to maximize the differentiation of articles, as the forensic community is relatively small and often uses similar language. Common language words and a set of forensic-specific words that did not add value to topic discovery (stopwords) were excluded from the text corpus to ensure they did not affect the analysis. Due to the multilingual nature of the data, multilingual stopwords from the package nltk (v3.9.4) were also used. A custom vectorizer model was included, as BERTopic only allowed stopwords to be removed if a custom vectorizer model was used.

#### Topic Representation

A topic representation model was used to improve the clustering of keywords because words such as *interpretation*, *Bayesian*, and *statistics* tended to dominate clusters without distinguishing between underlying topics. The KeyBertInspired function from the BERTopic Python module (v0.17.4) (Grootendorst, Maarten, 2025) was used to improve the representation of topics. This model uses an additional cluster based TF-IDF matrix (cTF-IDF) to support the clustering process based on random sampling of the records to find candidate keywords. Two thousand random samples (nr_samples) of 800 representative documents within each of the topics defined by the preceding steps (nr_repr_docs) were taken. Two thousand candidate words from the combined text corpus (nr_candidate_words) were assessed, returning the 20 most frequent words (top_n_words). These parameter values were chosen to provide robust sampling of terms and documents.

### Model Optimization

As BERTopic is a modular system with different components. It also has many different parameters that may be set. Combinations of different components and parameter values were investigated to determine the best model for analysis. The components adjusted were as follows and are shown in Figure 2 for clarity:

1. Use of BERTopic with no adjustments (default). This provides a baseline against which other combinations of components and parameters can be assessed.
2. Adding the custom Vectorizer model to the default BERTopic model (*Vectorizer only*). This allowed the addition of stopwords to better define the topics.
3. Using the ZeroShot candidate topics with the default inclusion similarity value of 0.7 (*ZS (De-fault) + Vectorizer*). It was hypothesized that by guiding the topics with a candidate list, more relevant topics would be found.
4. Adjusting the ZeroShot inclusion similarity to 0.8 (*ZS + Vectorizer*). Increasing the stringency of similarity required to be included in a topic should increase the coherence score for each topic.
5. Adjusting the default UMAP nearest neighbor parameter value to 20 for model 4 (*ZS + UMAP + Vectorizer*). This should guide the model to find a smaller number of larger topics, but it might reduce coherence.
6. Adding the cTF-IDF representation model to model 5 (*ZS + UMAP + Vectorizer + cTF-IDF*). This should improve the description of the topics without affecting the score.
7. Removing ZeroShot and changing the base Sentence Transformer model to paraphrase- multilingual-mpnet-base-v2 (*MLbase + Vectorizer*). Trying the multilingual model as the sole change from the Vectorizer only model was carried out to determine the effect of multiple languages being present in the dataset on topic determination.
8. Using both ZeroShot and the changed Sentence Transformer model with CountVectorizer (*ZS + MLbase+Vectorizer*). ZeroShot was added again to try to increase the relevance of the topics returned.
9. Adjusting the default UMAP nearest neighbor parameter value to 20 for model 7 (*MLbase + Vectorizer + UMAP*). This model was run to compare with the ZS + UMAP + Vectorizer model.
10. Adding a customized prompt to the Sentence Transformer model (*SBERT + Vectorizer*). Using a custom prompt allowed the model to be given context on the task. it was anticipated that this would result in clearer and more relevant topics.
11. Adjusting the UMAP nearest neighbor value to 20 for model 10 (*SBERT + Vectorizer + UMAP*). This model was run to compare with the ZS + UMAP + Vectorizer and the MLbase + Vectorizer + UMAP models.
12. Adding the cTF-IDF representation model to model 11 (*SBERT + Vectorizer + UMAP + cTF-IDF*).

**Figure 2:**
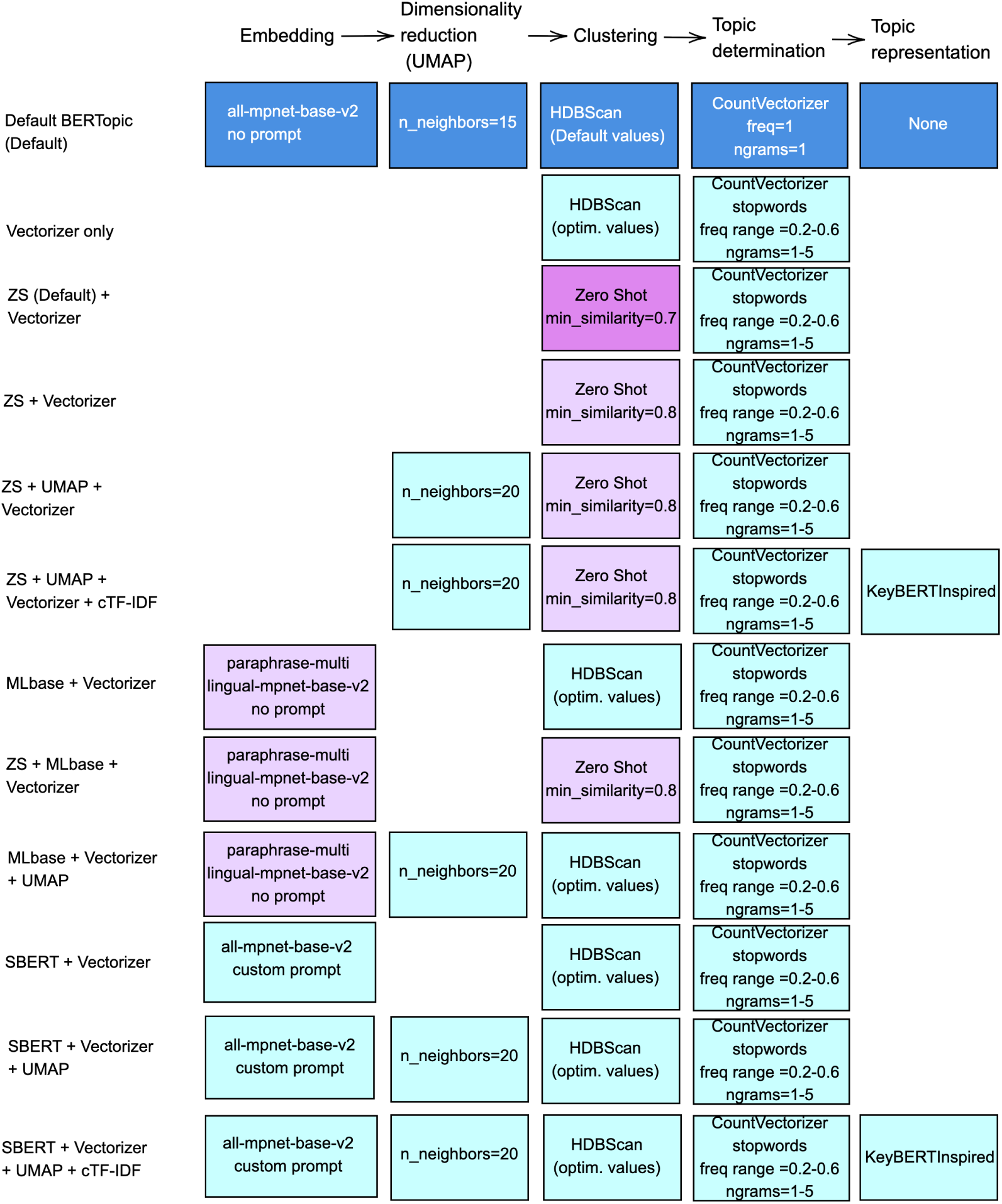
Altered parameters tested for optimization of BERTopic model. Colors used to indicated changed values.

Additionally, a system grid-search over the parameter space was used to optimize the hyperparameters for the HDBScan function for each of the above iterative changes. The only exception to this was for model 1, where the default cluster model parameters were maintained. Specifically, the ranges for minimum samples for cluster inclusion (a range of 5–25 by 5), minimum distance to the nearest neighbors (*k* parameter with a range of 1–10), and cluster inclusion threshold (*ɛ* parameter with a range of 0–0.4 by 0.1) were assessed.

### Model Comparison and Validation

The best model and parameters were chosen based on a combination of three metrics: topic coherence, diversity, and silhouette coefficient. These metrics are described below. A composite metric was decided upon as there was no clear consensus on the best approach in the literature. This overall composite score was weighted towards coherence, as shown in Equation 1, because obtaining coherent topics was the most desirable outcome.

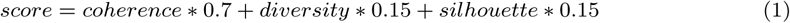

#### Topic Coherence

Topic coherence was calculated using the CoherenceModel function from the Python package Gensim (v4.4.0), with coherence = "c_npmi" to specify the *Normalized Pointwise Mutual Information* (NPMI) coherence measure (Bouma, 2009). This measure assesses the association between terms within a topic based on their co-occurrence in the original documents. NPMI values range from -1 to 1, with values closer to 1 indicating greater topic coherence and values closer to -1 indicating lower topic coherence. NPMI coherence is defined as

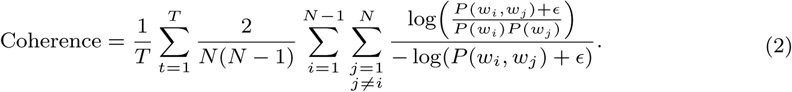

where

*• T* is the number of topics;
*• N* is the number of terms considered for each topic;
*• P* (*w_i_*) is the probability of observing term (or word) *w_i_* in the corpus;
*• P* (*w_i_, w_j_*) is the probability that terms *w_i_* and *w_j_* co-occur within the same document;
*• w_i_* and *w_j_* denote the *i*th and *j*th terms within topic *t*, where *i* ≠ *j*;
*• ɛ* is a smoothing constant of 10^−5^ to ensure that calculation errors do not occur (Röder et al., 2015).

#### Topic Diversity

Topic diversity, or simply diversity, measures the extent to which the topics identified by the model are distinct from one another.. A metric using a type-token ratio (TTR) approach -called the *Measure of Textual Lexical Diversity* (MTLD) (McCarthy, 2005) -was used in this study. The MTLD metric was chosen because it is tolerant of long or differing lengths of the text corpus being assessed while maintaining robust sensitivity to differences between documents (Torruella and Capsada, 2013). Specifically, the Python implementation of this metric from the package lexicalrichness (v0.5.1) was used. The lexicalRichness function combined all terms found for each topic and calculated a moving average sequential TTR value with a sliding window of one term. The calculation of TTR is shown in Equation 3, where *V* is the set of words in a topic and *N* is the total number of words for the topic. The number of text fragments for which this value remained above a threshold of 0.72 was recorded in both forward and reverse reading directions and then averaged. A higher average number of fragments indicates more distinct topics.

where

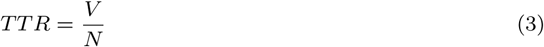

*• V* is the number of unique words in the topic;
*• N* is the total number of words in the topic.

#### Silhouette Coefficient

The Silhouette coefficient (Rousseeuw, 1987) was calculated using the silhouette_score function from the Scikit-learn (v1.9.0) module. The Silhouette coefficient determines the average dissimilarity of each document to all other documents within the same topic and compares it to documents in the nearest neighboring topic to which it could otherwise be conceivably assigned. See Equation 4, where *N* is the set of documents, *C_A_*is the topic in which the document is classified, *C_B_* is a different topic, and *d*(*i, j*) is the euclidean distance between *i* and *j*.

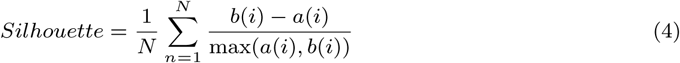

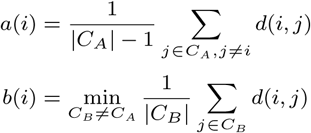

where

*• N* is the number of documents;
*• i* denotes the document for which the silhouette value is being calculated;
*• j* denotes another document used in the distance calculation;
*• C_A_* is the topic to which document *i* is assigned;
*• C_B_* is a topic other than *C_A_*;
*•* |*C_A_*| and |*C_B_*| are the numbers of documents assigned to topics *C_A_* and *C_B_*, respectively;
*• d*(*i, j*) is the Euclidean distance between documents *i* and *j*;
*• a*(*i*) is the mean distance between document *i* and all other documents assigned to the same topic;
*• b*(*i*) is the smallest mean distance between document *i* and the documents assigned to any other topic.

After the parameters of the best scoring model were selected, the model was then assessed for robustness. This assessment was conducted using repeated five-fold cross-validation with the number of repeats set to five. This was implemented using the RepeatedKFold function from the scikit learn model_selection module. The parameters for HDBScan and the number of topics were set manually to ensure consistency. Cross-validation examined how stable the topics were across different fittings of the data, the stability of the scoring, and the mean number of outliers that were detected. Ninety-five percent confidence intervals (95% CI) were calculated using Student’s *t*-distribution. The appropriate number of topics was also investigated, with the overall score being optimized across a set number of topics between 10 and 36.

### Visualization

Clusters were visualized as an *inter-topic distance map* using BERTopic’s built-in UMAP, with Euclidean distance as the determining metric. Custom cluster labels were added to ensure clarity. Hierarchical clustering was carried out to visually explore the relationships between the topics.

## Results

### Investigation of the viability of automated literature search

#### LiteratureReview tool

Use of the LiteratureReview tool requires knowledge of the Python programming language and the ability to run Python code from the command line. Some example code is provided with the tool, but inexperienced or first time Python users may struggle. The tool uses Selenium, a common tool for obtaining information from websites. An issue with the library is that it acts like a human interacting with a website, including clicking buttons and filling in fields. If the content of the website changes, it will break down. In the two years since the code (Hewarathna A. I., 2023) was written, the databases accessed have altered the allowable features of their systems to reduce the ability of computers to automatically obtain the data. Features such as checking for human interaction, requiring login credentials, changing where items are located on the page, how they are named, and initiating terms and conditions for the use of their search tools to exclude AI are now in place. Additionally, the codebase for the tool was last updated in 2023, indicating that it is likely not maintained by the developer. For these reasons, this tool is not a viable long-term solution for use, especially not for a forensic use-case where findings can be interrogated in a court of law. No data was added to the analysis from this tool.

#### AI Scientist and Semantic Scholar

The installation and use of the AI Scientist tool are well-documented, but it requires users to be familiar with command-line interaction with Python, specifically the Anaconda distribution and environment. The majority of the code included in the AI Scientist repository was not relevant to a forensic literature recovery task, as it was designed for computer science research. At the time of this investigation, this tool also did not have access to forensic-specific journals where TPPR data are most likely to be published. Therefore, the implementation of this component of the AI Scientist tool did not progress further. This issue was monitored during the course of this investigation for improvements to the Semantic Scholar results, as this API is a legal and viable way to obtain information relating to forensic papers, if they are made available. During the course of this investigation, the relevance of articles improved; however, there was still a lag in available papers of up to three years. No results were added to the analysis from this tool.

#### Clarivate API

The search query returned 889 unique records, each containing 38 metadata fields, although many of these fields were incomplete or missing. These records were combined with those from the forensic databases for further analysis. Of these 889 records, 680 were not present in the forensic databases. While abstracts were present in the data, the experimental data factors relevant to Bayesian Network (BN) development were not, nor were any supplemental materials obtained through this means. However, a comparison of the results to other sources provided increased article metadata and additional articles that were missed or excluded from forensic repositories.

#### Semantic Scholar web interface

The initial use of the Web of Science query 1 did not work, with Semantic Scholar unable to understand such a complex variety of conditions. The query was altered to ”forensic evidence transfer persistence prevalence recovery“ which resulted in a return of 574 results. These appeared mostly relevant to the query, although some forensic psychology, economic and sociologically related papers were observed. The web interface does not offer the ability to download the results nor filter the results using keywords. Visual inspection of the results suggested that most of the records recovered were already in the other data sources.

### Find and access existing data sources

#### Organizational repositories

The organizational repositories comprised three complementary collections: the University of Lausanne repository containing 77 journal articles and 11 theses; 58 TPPR publications identified during a literature review (Stacey et al., 2025); and a bibliography of 751 records compiled by Dr. John Butler. These repositories contributed 398 records that were not identified from any other source, comprising 16 publications from the internal repositories and 382 from Dr. Butler’s bibliography.

#### DNA-TrAc

DNA-TrAc contained 311 records described by 44 metadata fields, together with a supplementary list of publications awaiting incorporation into the database. Twenty-eight of these records were not present in the TTADB.

#### Transfer Traces at Activity Level Database (TTADB)

The TTADB contained 2438 records described by 15 metadata fields. The collection included publications in multiple languages, including English, French, Polish, and German.

### BERTopic Model Optimization

Combining all datasets resulted in a single dataframe with 3279 unique records and 88 fields. Fortyseven (47) of these fields were concatenated for analysis. These are listed in B. The full list of records is available on request. BERTopic model optimization results are shown in Table 2. The aim of optimization was to maximize all scores, with coherence and silhouette having a maximum value of 1, and diversity and the overall score having a maximum of infinity.

**Table 2.** BERTopic model optimization results. HDBSCAN values are reported in the order minimum cluster size, minimum samples, and cluster-selection epsilon. Models marked NA did not include an HDBSCAN clustering step. Overall scores were calculated using the unrounded component scores. Overall Score was calculated as per Equation 1.

| **Model** | **Model Components** | **HDBSCAN Values** | **Coherence Score** | **Diversity Score** | **Silhouette Score** | **Overall Score** | **Number of Topics** |
|---|---|---|---|---|---|---|---|
| 1 | Default | 10, 0, 0.0 | -0.18 | 80 | 0.064 | 12 | 19 |
| 2 | Vectorizer only | 5, 5, 0.1 | -0.17 | 420 | 0.004 | 62 | 37 |
| 3 | ZS (Default) + Vectorizer | NA | -0.12 | 320 | 0.035 | 47 | 47 |
| 4 | ZS + Vectorizer | NA | -0.20 | 420 | 0.018 | 63 | 32 |
| 5 | ZS + UMAP + Vectorizer | NA | -0.33 | 310 | 0.064 | 46 | 21 |
| 6 | ZS + UMAP + Vectorizer + cTF-IDF | NA | -0.16 | 100 | -0.037 | 14 | 21 |
| 7 | MLbase + Vectorizer | 5, 2, 0.2 | -0.15 | 420 | 0.014 | 63 | 33 |
| 8 | ZS + MLbase + Vectorizer | 5, 1, 0.2 | -0.24 | 450 | 0.059 | 67 | 28 |
| 9 | MLbase + Vectorizer + UMAP | 5, 5, 0.1 | -0.13 | 380 | 0.038 | 57 | 35 |
| 10 | SBERT + Vectorizer | 5, 4, 0.2 | -0.15 | 410 | 0.020 | 62 | 28 |
| 11 | SBERT + Vectorizer + UMAP | 5, 1, 0.1 | -0.15 | 320 | 0.018 | 48 | 46 |
| 12 | SBERT + Vectorizer + UMAP + cTF-IDF | 5, 1, 0.1 | -0.09 | 110 | 0.011 | 17 | 53 |

All of the models that were explored returned negative coherence scores, indicating that the documents within each topic were not particularly similar to one another. This is expected, as publications on different trace types explore many aspects of variation. Model 12 returned the best performing coherence score, indicating that the topics were the most distinct and logical out of the models tested. However, the coherence score was still negative, indicating that this was not highly coherent. The slight gain in coherence was also offset by a large loss in diversity compared to other models. Diversity scores were high across most of the model iterations, suggesting that articles assigned to different topics consistently used different language. The addition of changes in the UMAP dimensionality reduction parameters or the use of a cTF-IDF representation model did not improve any model, with all scoring metrics showing a decrease with their use.

HDBScan parameter values remained largely static across the different models, with the minimum cluster size changing from the default value of 10 to a value of 5 in all models where clustering was performed. The minimum number of samples in a cluster and the cluster inclusion threshold values also remained low. As the minimum cluster size or minimum number of samples for inclusion increased, the reduction of the diversity score indicated that common language began to be more represented in the topics. This is likely due to the wide ranging aspects of investigation in studies.

Models that used ZeroShot classification often did not use HDBScan for clustering, so changing these parameters was not relevant.

Five of the models (Models 2, 4, 7, 8, and 10) showed a clear lead in performance, and all were comparable in overall score. All used a custom vectorizer model, regardless of the embedding method employed. The increase in the overall score was predominantly driven by the diversity score, with all of these models returning diversity scores exceeding 400. Model 8 was the highest performing model overall, with a combined score of 67. This score was obtained after an investigation was carried out to determine the most optimal number of topics when using the Model 8 parameters. This investigation is explained below.

Hierarchical clustering (Figure 3) provided a better view of the relationships between topics. It showed an early split between forensic biology and other traces. The clusters described appeared reasonable and could broadly be categorized into the following groups:

*•* explosives;
*•* hairs, fibers, GSR, paint, glass, ignitable liquids, illicit drugs, and soil;
*•* periand postmortem;
*•* fingerprints;
*•* DNA typing;
*•* trace DNA and transfer;
*•* bloodstain pattern analysis (BPA) and body fluid identification;
*•* sexual assault.

**Figure 3:**
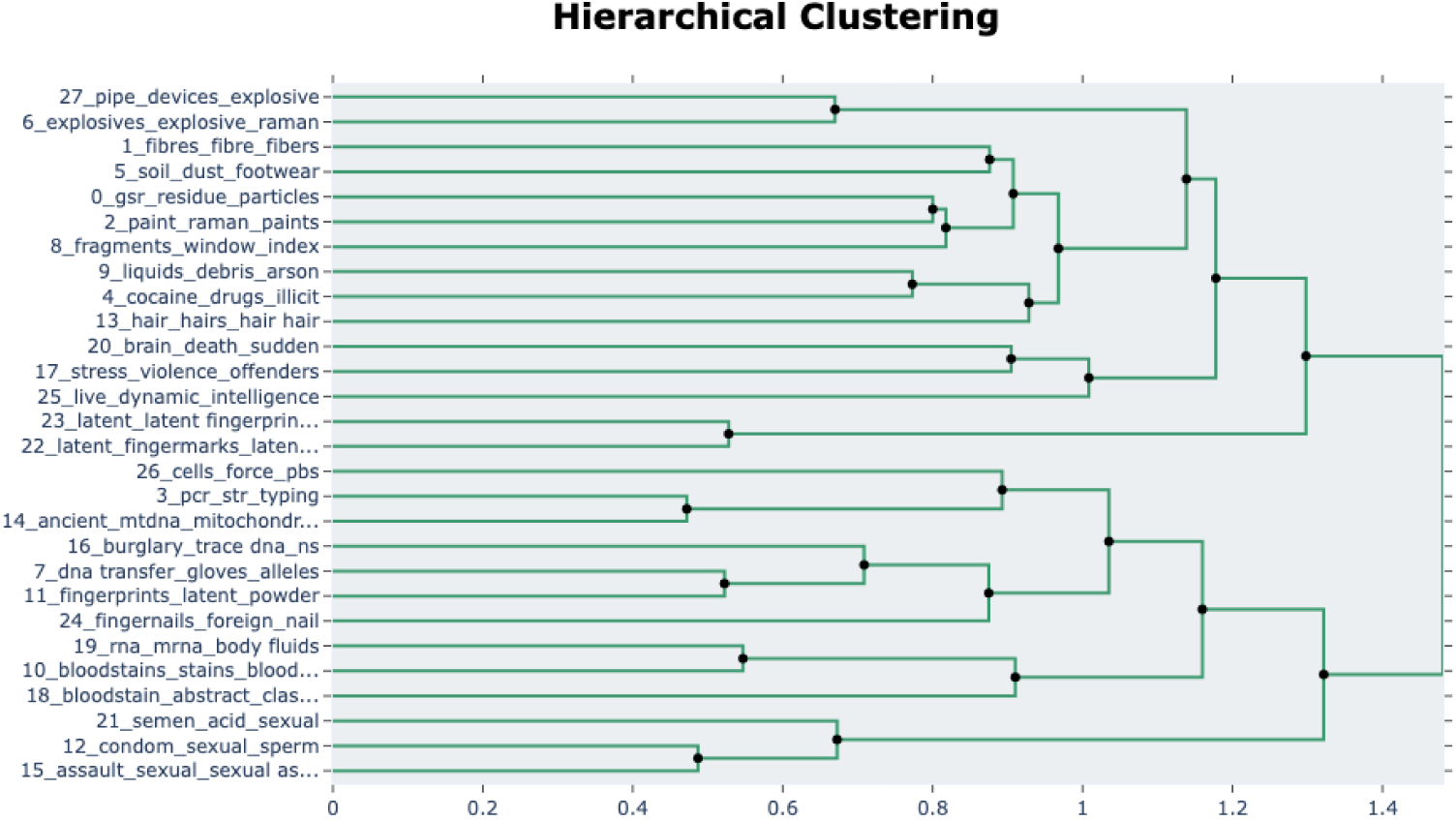
Hierarchical clustering of all topics. The X-axis shows a distance function based on 1 -cosine similarity of the topics based on their terms

This hierarchical view showed that the close alignment of some topics, such as topics 15 and 17, was not as significant as it appeared in the distance map. This may be due to the distance map being viewable only in two dimensions. This clustering method also demonstrated that the number of topics could be adjusted to simplify the topics by combining closely related topics. This was investigated to determine the most appropriate number of topics. Scoring was assessed when the number of topics was set between 10 and 36. The optimal number of topics based on the highest achievable score was assessed as 28. The scoring information for this version of the model is presented above.

The highest performing model (Model 8) was tested for robustness using repeated five-fold cross-validation. Overall, the terms describing the topics remained quite constant across the folds, with a mean cosine similarity value of 0.86. Interestingly, the final topic almost always described a single study on pipe bombs, indicating that this paper (Oxley et al., 2001) deviated from the language commonly used in studies on explosive materials. The mean number of outlier data points across the five folds was 22.3% (95% CI 20.5% − 24.1%). Cross-validation splits the data into five subsets, using four for training the model and the fifth for testing. Due to the randomized sampling used to decide the data subsets, the calculation of coherence for each fold was affected. The reduced input data caused some folds to contain either zero or only a single topic. A placeholder value of zero was used to ensure that the overall score calculation could proceed with no input from the coherence. When coherence could be calculated, it returned a positive value, with a mean of 0.063 (95% CI 0.047 − 0.078). The mean diversity of the folds was 238.6 (95% CI 197.1 − 280.1), and the mean Silhouette score was 0.075 (95% CI 0.071 − 0.079). The mean overall score obtained for the folds was 35.8 (95% CI 29.6 − 42.1). The reduction in performance from the initial high score obtained is likely due to the reduced data set that was assessed in cross-validation. The high cosine similarity of the topics output from each cross-validation fold, along with the reasonable confidence intervals of each scoring metric, indicates that the model is stable over multiple executions and is fit for use.

The topics obtained from Model 8 (ZS + MLbase + Vectorizer) were investigated further. The model returned 28 unique topics, and 703 of the 3279 documents were labeled as outliers, having support for inclusion lower than the similarity threshold for all topics. The majority of these outlier documents were related to DNA. This is a reasonably high proportion, and mechanisms exist to reduce this number by half (363). However, carrying out this outlier reduction caused the topics to become less defined, so it was not continued. The remaining topics are shown in Table 3. Topic 0 was the largest topic, with 583 records on gunshot residue (gsr). Forensic biology related records were split over 10 different topics.

**Table 3.** Topics determined using the ZeroShot+MLbase+Vectorizer model with number of documents assigned to each topic

| Topic | Name | Count |
|---|---|---|
| 0 | gsr_residue_particles_firearm | 583 |
| 1 | fibres_fibre_fibers_fiber | 215 |
| 2 | paint_raman_paints_spectroscopy | 194 |
| 3 | pcr_str_typing_dna dna | 191 |
| 4 | cocaine_drugs_illicit_hair | 151 |
| 5 | soil_dust_footwear_soils | 148 |
| 6 | explosives_explosive_raman_residues | 119 |
| 7 | dna transfer_gloves_alleles_substrate | 113 |
| 8 | fragments_window_index_broken | 111 |
| 9 | liquids_debris_arson_carpet | 94 |
| 10 | bloodstains_stains_bloodstain_rna | 90 |
| 11 | fingerprints_latent_powder_fingermarks | 74 |
| 12 | condom_sexual_sperm_vaginal | 74 |
| 13 | hair_hairs_hair hair_human hair | 68 |
| 14 | ancient_mtdna_mitochondrial_mitochondrial dna | 55 |
| 15 | assault_sexual_sexual assault_children | 50 |
| 16 | burglary_trace dna_ns_success rates | 39 |
| 17 | stress_violence_offenders_services | 38 |
| 18 | bloodstain_abstract_classification_discussing | 29 |
| 19 | rna_mrna_body fluids_body fluid | 22 |
| 20 | brain_death_sudden_injury | 21 |
| 21 | semen_acid_sexual_stains | 20 |
| 22 | latent_fingermarks_latent fingerprints_fingerprints | 19 |
| 23 | latent_latent fingerprint_luminescence_latent fingerprints | 17 |
| 24 | fingernails_foreign_nail_exogenous | 15 |
| 25 | live_dynamic_intelligence_framework | 14 |
| 26 | cells_force_pbs_sterile | 11 |
| 27 | pipe_devices_explosive_ | 1 |

The descriptive terms used to name the topics illustrate that the records were grouped into largely reasonable clusters. Some of the chosen terms representing smaller clusters may be too broad for an ALE use case, although this may depend on the question of interest. For example, “cells_force_pbs_-sterile”. Additionally, although the search query used with the Web of Science API specified filtering of digital forensics, forensic psychology, and astrophysics, some studies remained in the dataset. Some of these records were proposed as topics, such as “live_dynamic_intelligence_framework” encompassing digital traces and “stress_violence_offenders_services” containing psychology studies, while others were correctly removed as outliers. Records related to firearms and gunshot residue comprised the largest topic (Topic 0) with 583 records.

When the topics shown in Figure 3 were visualized with UMAP, some differentiation was observable. An inter-topic distance map is shown in Figure 4. Some clustering can be detected, with topics 0 (gunshot residue), 6 (explosive residues), 9 (ignitable liquids), and 27 (pipe bomb) creating a well-defined cluster. Topics 14 (sexual assault), 15 (sperm and sexual intercourse), 17 (violence and offender services), 20 (condoms), and 24 (semen identification) were, somewhat unsurprisingly, closely clustered and quite distinct from other topics. Interestingly, topic 2 (paint) clustered with topics 10 (presumptive testing of bloodstains), 18 (RNA body fluid identification), 19 (BPA), and 20 (death and postmortem injuries).

**Figure 4:**
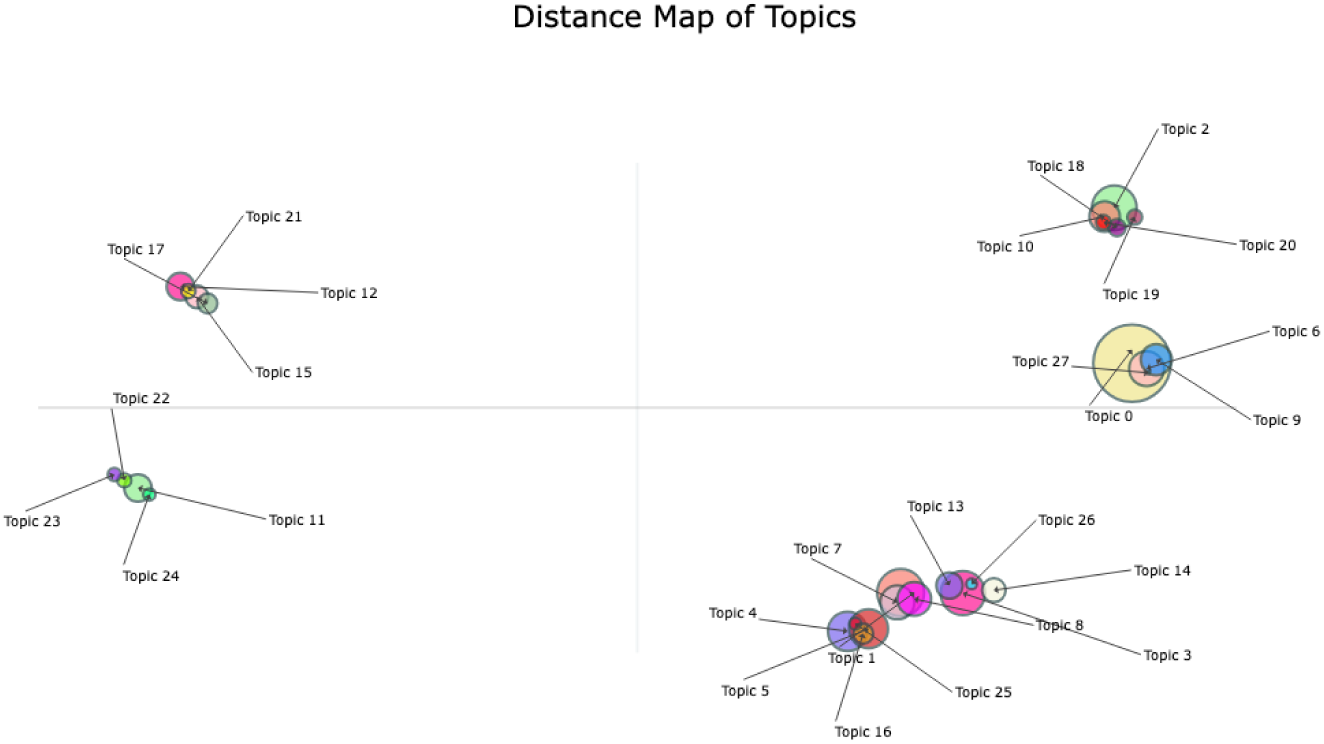
Inter-topic Distance Map of all data records. The size of the topic circle is proportional to the number of records classified in the topic.

Topics 12 (condoms), 15 (sexual assault), 17 (stress and offender services), and 21 (semen stains) somewhat unsurprisingly formed a cluster. Equally, it would not be considered surprising that Topics 11 (fingerprint enhancement), 22 (latent fingerprints), 23 (fingerprint visualization), and 24 (nails and foreign DNA) form a distinct cluster.

The remaining topics appeared to group into a single elongated cluster, but a closer examination showed three sub-clusters. The first comprised topics 4 (illicit drugs), 5 (soil, dust, and plant material), 16 (trace DNA), and 25 (digital traces). Topics 1 (fibers), 7 (DNA transfer), and 8 (glass) comprised the second cluster, and the third cluster was made up of topics 3 (DNA typing), 13 (hairs), 14 (extracellular DNA collection), and 26 (mitochondrial DNA).

## Discussion

This study investigated whether automated literature retrieval and natural language processing (NLP) methods could support the identification, organization, and characterization of TPPR literature relevant to forensic activity level evaluations (ALE). The findings demonstrate that while automated literature retrieval is possible, further work needs to be carried out to support its relevance for a forensic use case. The evaluation of automated literature retrieval tools revealed substantial variation in their suitability for forensic applications. Tools based on web scraping, such as LiteratureReview, were found to be fragile and highly dependent on website structure, making them unsuitable for long-term forensic use where reproducibility and transparency are critical. Similarly, although the AI Scientist framework offered promising capabilities, their fundamental development use case in computer science and limited coverage of forensic journals restricted their usefulness for the current application. Additionally, the landscape of automated tools is fast-moving, and better ways of organizing access to literature are constantly being developed. Semantic Scholar initially showed a lack of forensic articles; although this improved over the course of the study. Manual access to the studies was difficult, but using the API may be a viable access route in the future. A lag in the indexing of newly published studies by Semantic Scholar may still pose a risk that evaluations carried out may not include the most up to date knowledge. Direct access to both the Semantic Scholar and ScienceDirect databases through an official API would be a stable, legally compliant, and transparent method of accessing records. However, while both are capable of returning a large number of records, not all of these are relevant, and both of these APIs only provide abstracts rather than full texts. These findings suggest that future automated forensic literature retrieval systems may benefit from a combination of multiple methods and, preferably, full-text access permissions.

The integration of multiple repositories and automated retrieval through the Web of Science API in this study resulted in the assembly of a corpus comprising 3,279 unique records. This suggests that concerns regarding a complete lack of supporting literature for activity level evaluations may be overstated. Multiple studies were identified for several trace types, including gunshot residue, fibers, paint, explosives, trace DNA, bloodstains, fingerprints, and illicit drugs.

Topic modeling, when carried out on a given dataset, could be a feasible route for filtering records for specific topics. While the use of BERTopic models has been superseded by larger, more powerful public models, this methodology was considered an analytical option because it offered benefits for a forensic science use case. The component-based nature of BERTopic provided transparency into how decisions were made; it allowed customization at a granular level to ensure the accuracy of information; and the size of the Sentence transformer and ZeroShot classification models was small enough to run locally, thus ensuring data protection and security. Furthermore, the results highlight both the considerable breadth of published TPPR research and the challenges associated with making that information readily accessible and suitable for evaluative use.

The BERTopic modeling approach successfully identified meaningful thematic groupings across the combined dataset, even though the model was not given full texts. Despite relatively low coherence scores, the resulting topics largely reflected recognizable forensic disciplines and trace evidence categories. Major topic clusters corresponded to gunshot residue, fibers, paint, explosives, soil, trace DNA, blood evidence, body fluid identification, fingerprints, and sexual assault investigations. Importantly, these topics emerged without manual categorization of individual studies, suggesting that transformer-based NLP methods can identify meaningful relationships within heterogeneous forensic literature. Seven hundred and three records were considered outliers. Many of these were DNA related, indicating there may be further underlying topic structure that was not characterized and requires further exploration.

The relatively poor coherence values observed across all model variants warrant consideration. While low coherence scores would normally suggest weak topic definition, they may be expected in this context. TPPR studies investigate a wide range of experimental variables, trace materials, transfer mechanisms, environmental influences, and analytical methods. As a result, documents grouped within the same broad forensic topic may still differ substantially in language and experimental focus. Consequently, conventional topic-modeling performance metrics developed for more homogeneous corpora may underestimate topic quality in forensic datasets. This interpretation is supported by the generally sensible topics produced and the high topic stability observed during cross-validation. Additionally, the low coherence scores may be an artifact of the small corpus of text available per document. Analysis of full-texts may improve this metric.

Interestingly, diversity scores were consistently high across most model configurations. This indicates that different topics employed substantially different vocabularies, suggesting that the literature contains distinguishable disciplinary domains. The strong performance of models incorporating custom vectorization further emphasizes the importance of domain-specific preprocessing. The removal of generic forensic terminology and multilingual stopwords appeared to improve topic separation more effectively than modifications to dimensional reduction or representation models.

The optimized model also demonstrated good robustness. Topic labels remained highly consistent across repeated cross-validation runs, with a mean cosine similarity of 0.857. This stability suggests that the identified themes are not artifacts of random initialization or specific document subsets but instead reflect genuine structure within the literature. Although approximately 22% of records were classified as outliers, many of these were related to DNA that did not align strongly with the candidate topics. Reduction of outliers might be achieved by re-investigating the chosen candidate topics for ZeroShot Classification or by subsetting forensic biology related studies and performing topic modeling within the discipline.

Combining data from current forensic knowledge repositories with results from a standard literature database search provides increased information. DNA-TrAc contains highly detailed experimental descriptors specifically relevant to forensic biology studies, while the TTADB offers broader coverage across trace evidence disciplines but contains less detailed experimental metadata. These differences reflect the distinct purposes, scope, and technical characteristics of the available resources rather than their overall quality. The addition of data from the Web of Science and CrossRef APIs enriched both datasets, improving article identification and providing access to records absent from existing forensic databases. These findings support the value of integrating multiple sources rather than relying on any single repository.

Several observations have important implications for activity level evaluations. First, the large number of studies identified within some trace evidence domains suggests that opportunities exist for more evidence-based approaches to evaluating activity level propositions. Second, topic modeling may provide a practical method for rapidly identifying literature relevant to a particular trace type or scenario. Such functionality could reduce the burden on practitioners searching for supporting evidence and improve awareness of newly published research. Third, the integration of repository data with automated retrieval systems could form the basis of a continually updated forensic knowledge base capable of supporting evaluative reporting and Bayesian network development.

However, a large volume of literature does not automatically imply suitability for ALE. This study evaluated literature availability rather than evidential quality, and the primary challenge appears to be less the absence of studies and more their discovery, assessment, and reuse. Recent inter-laboratory studies have demonstrated substantial variability in TPPR findings (Gill et al., 2025), while many publications lack sufficient methodological detail for direct comparison or incorporation into evaluative frameworks (Gill et al., 2026). Future work should therefore focus on assessing study quality, experimental validity, transferability, and case-specific relevance.

Several limitations should be acknowledged. The corpus used in this study was restricted to selected repositories and databases; while broad, it does not represent all published TPPR literature. It has been constrained by the existence of publisher paywalls and is not inclusive of studies in all languages (although the model would cope with additional languages). It should also be noted that books contain more information than journal articles and may give a false sense of the amount of data available for some topics. Topic modeling performance was influenced by preprocessing decisions, candidate topic selection, and model configuration. Furthermore, the analysis relied primarily on abstracts, metadata, and repository summaries rather than full-text articles, limiting the use of information contained in methods, supplementary materials, and experimental results. Future studies incorporating full-text analysis and leveraging citation networks, retrieval-augmented large language models, and structured metadata extraction may provide a more detailed characterization of the TPPR data landscape.

Until a correction method can address inter-laboratory variability, published results may provide only a broad indication of expectations. Nevertheless, the substantial number of available studies suggests that ALE may be viable in some cases when applied with appropriate caution. Automated systems for the continuous identification, categorization, and updating of TPPR studies, combined with semantic search and quality assessment frameworks, could further increase the utility of the literature. The implementation of minimum reporting standards (Gill et al., 2026) and correction methods (Taylor et al., 2025) may also improve interoperability and reuse of published data across laboratories and jurisdictions.

In conclusion, this study demonstrates that NLP-based analysis of TPPR literature, based on abstracts alone, is both feasible and informative. By combining forensic repositories with automated literature retrieval and transformer-based topic modeling, a broad and structured view of the available studies was obtained. The findings suggest that substantial TPPR literature already exists across multiple forensic disciplines and that automated approaches can assist in its discovery and organization. While challenges remain regarding data quality, standardization, and practical applicability to activity level evaluation, the methods presented here represent a promising step toward the development of scalable, maintainable, data-driven knowledge systems for forensic science.

## Conflicts of interest

The authors declare that they have no competing interests.

## Funding

This work was carried out as part of a PhD in Forensic Science degree at The University of Auckland and was supported by the New Zealand Ministry of Business, Innovation & Employment Strategic Science Investment Fund.

## Ethics

All data used in this study were obtained from publicly available sources. No ethics approval was required for the completion of this work.

## Data availability

The data underlying this article are available upon request from the corresponding author.

## Author contributions statement

J.S. conceived the study, collected and curated the data, carried out the experiment, and drafted the manuscript. J.M.C and S.A.H provided supervision and resources. S.O and S.A.H. provided acquisition of funding and project administration. All authors edited and reviewed the manuscript.

## Supporting information

Interactive Topic Map

## Acknowledgments

The authors thank the anonymous reviewers for their valuable suggestions.

## Appendix 1 -Data Source Fields

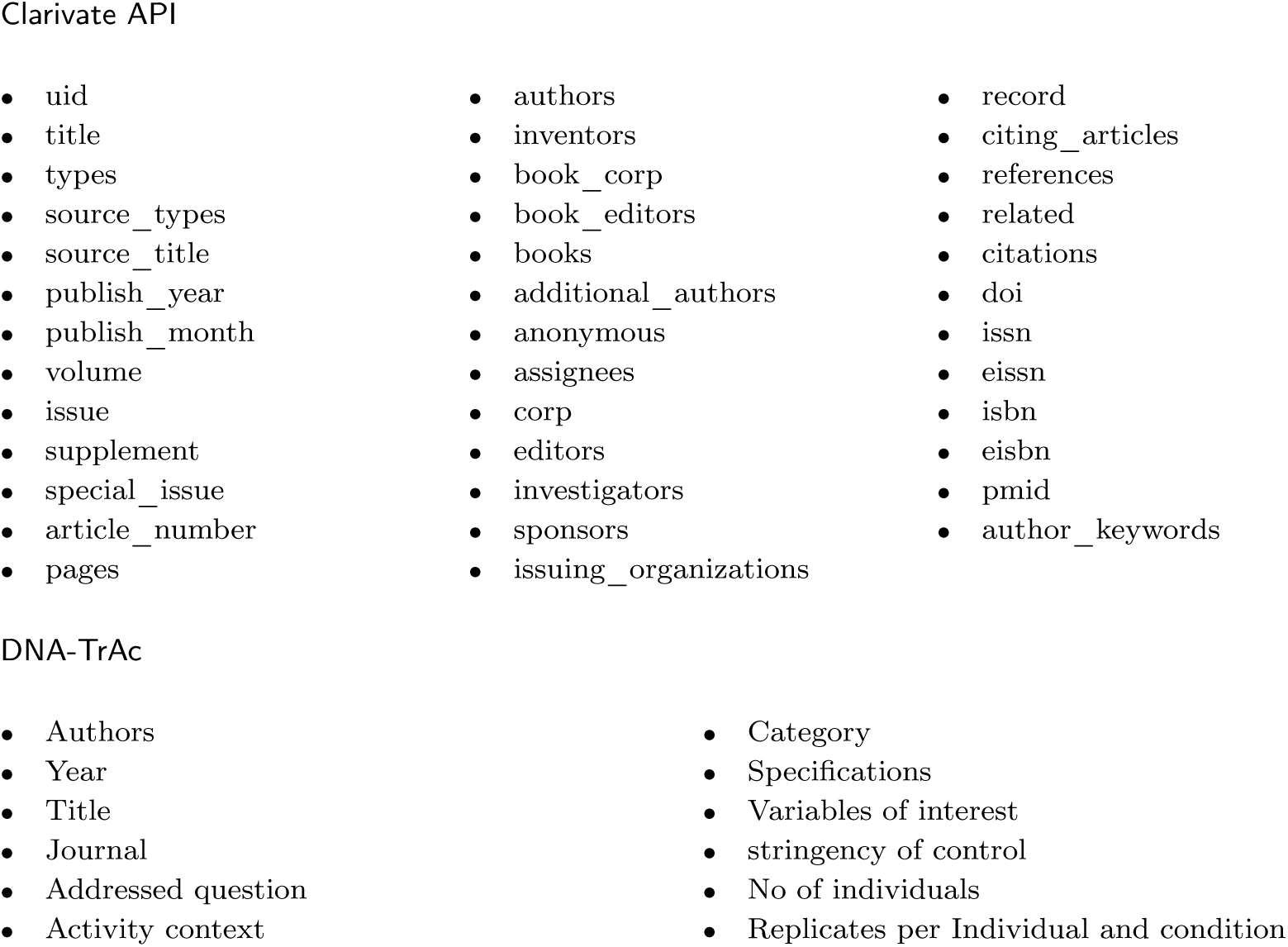
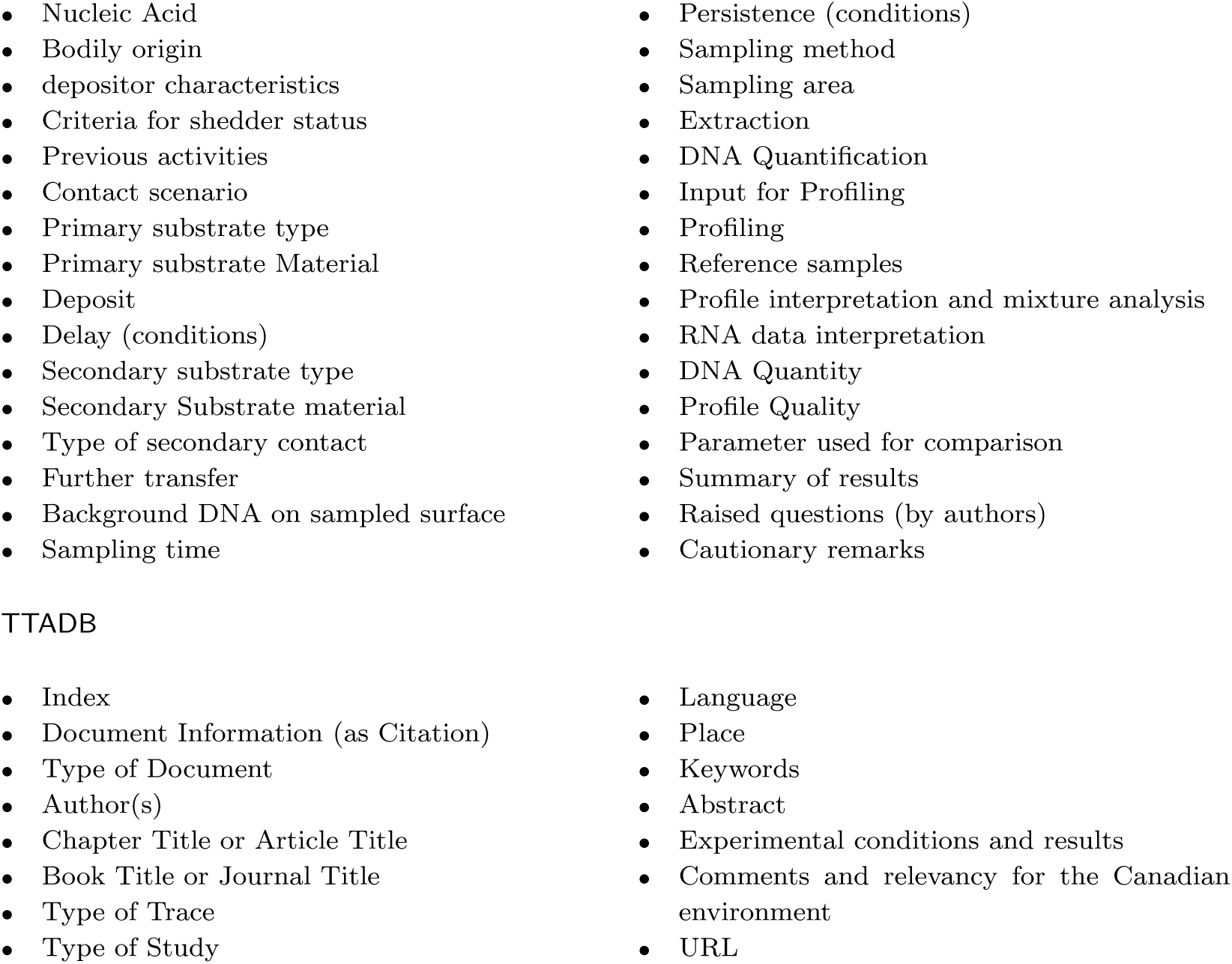

## Appendix 2 -Data Fields concatenated for analysis

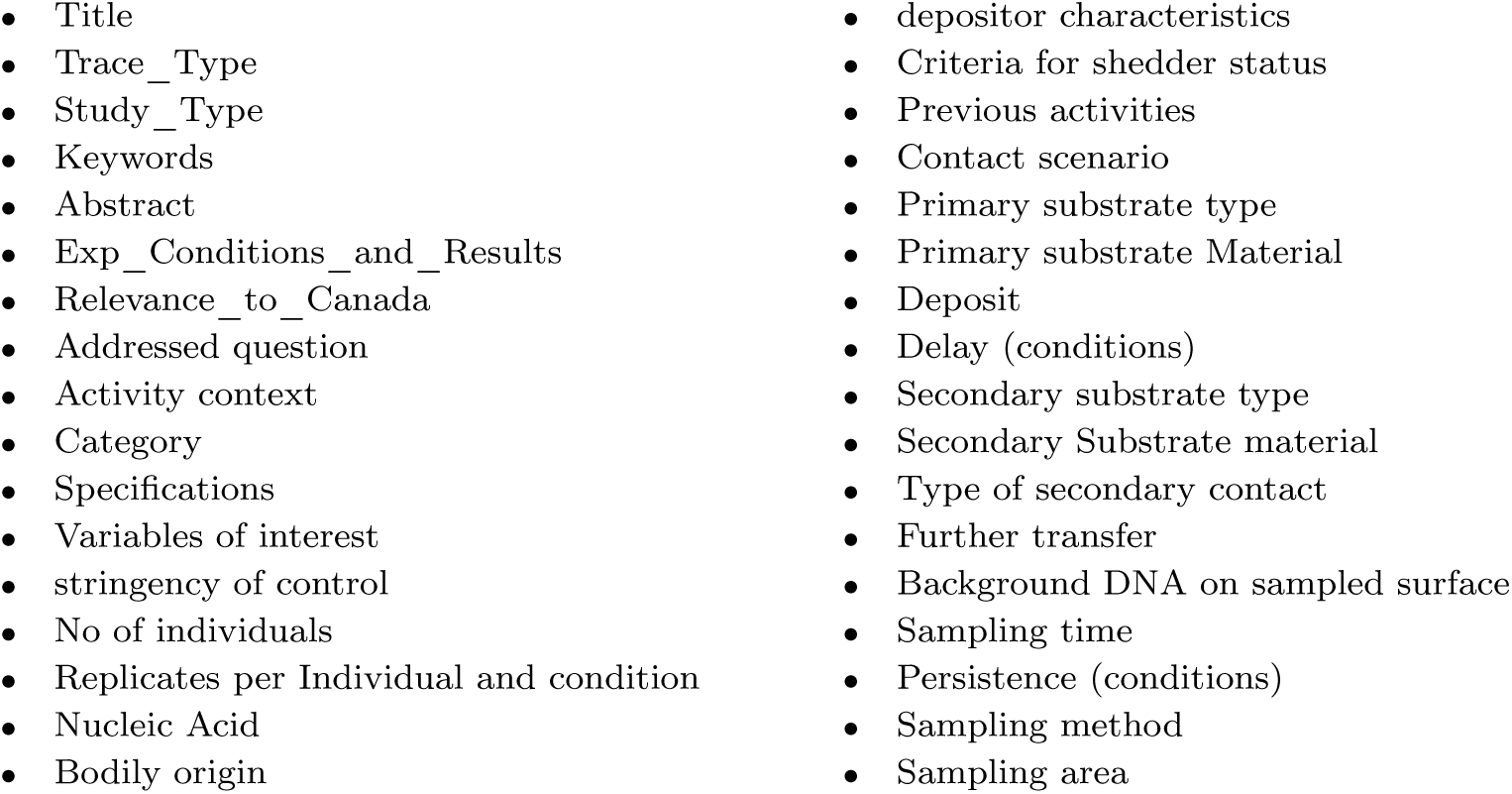
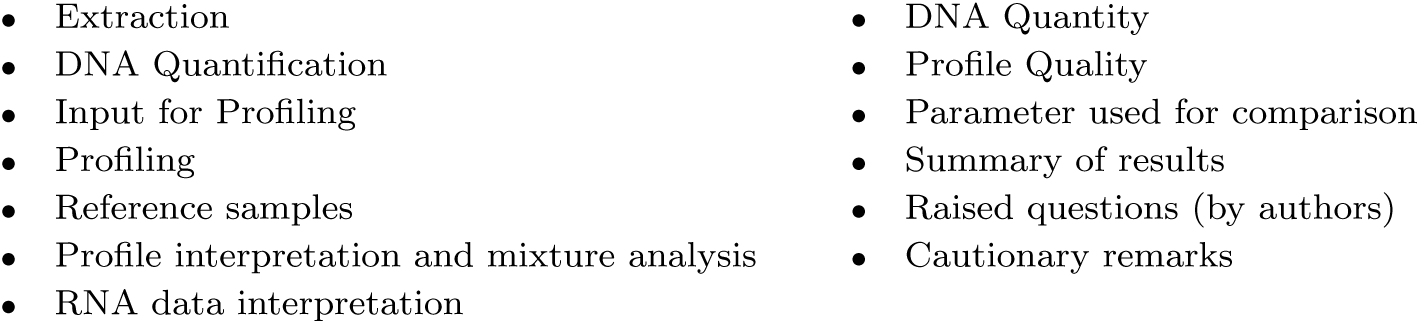

|  |  |
|---|---|
| Title | depositor characteristics |
| Trace_Type | Criteria for shedder status |
| Study_Type | Previous activities |
| Keywords | Contact scenario |
| Abstract | Primary substrate type |
| Exp_Conditions_and_Results | Primary substrate Material |
| Relevance_to_Canada | Deposit |
| Addressed question | Delay (conditions) |
| Activity context | Secondary substrate type |
| Category | Secondary Substrate material |
| Specifications | Type of secondary contact |
| Variables of interest | Further transfer |
| stringency of control | Background DNA on sampled surface |
| No of individuals | Sampling time |
| Replicates per Individual and condition | Persistence (conditions) |
| Nucleic Acid | Sampling method |
| Bodily origin | Sampling area |
| Extraction | DNA Quantity |
| DNA Quantification | Profile Quality |
| Input for Profiling | Parameter used for comparison |
| Profiling | Summary of results |
| Reference samples | Raised questions (by authors) |
| Profile interpretation and mixture analysis | Cautionary remarks |
| RNA data interpretation |  |

## Appendix 3 -Candidate Topic List for Zero Shot Classification

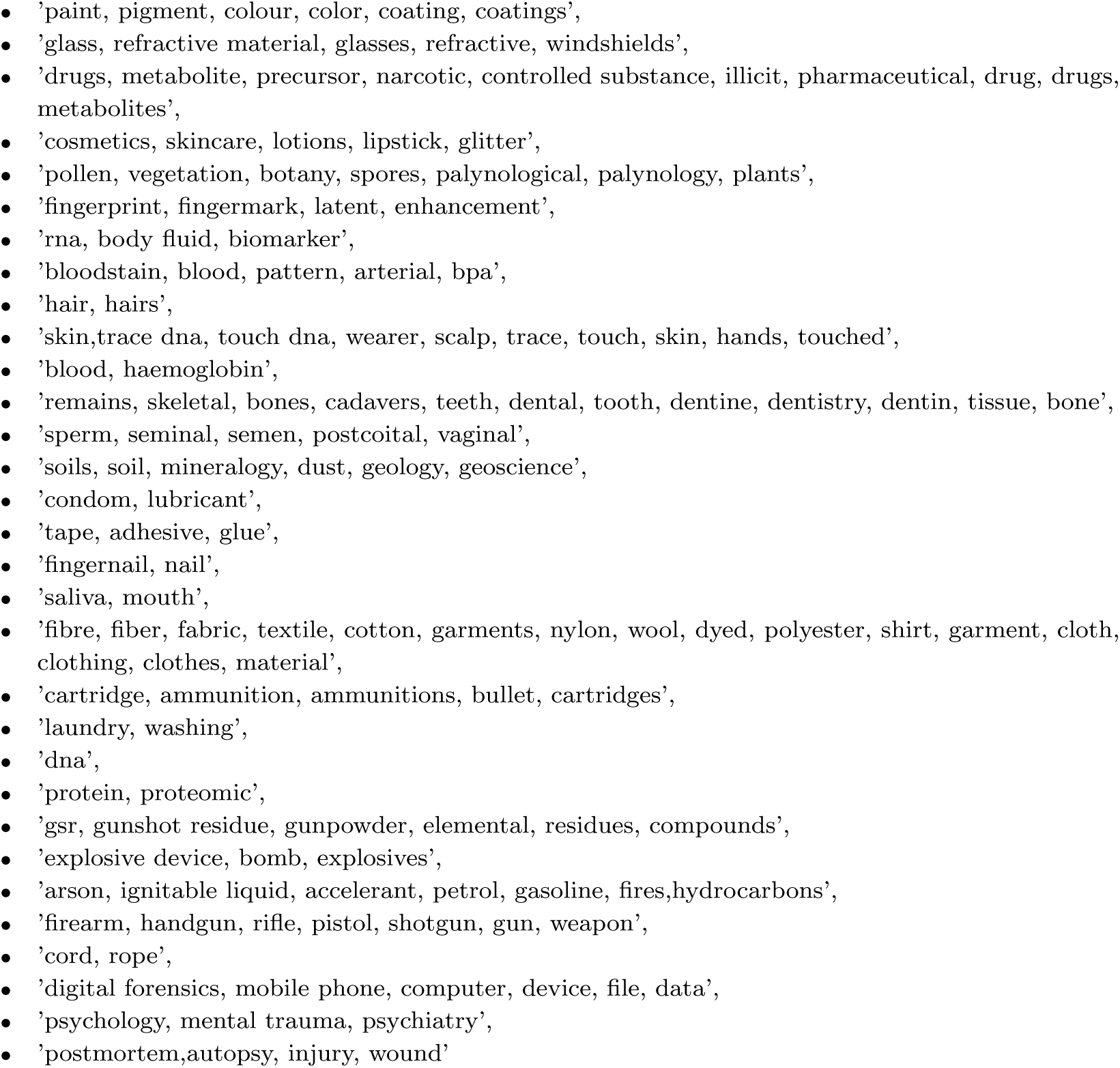

|  |
|---|
| ’paint, pigment, colour, color, coating, coatings’, |
| ’glass, refractive material, glasses, refractive, windshields’, |
| ’drugs, metabolite, precursor, narcotic, controlled substance, illicit, pharmaceutical, drug, drugs, metabolites’, |
| ’cosmetics, skincare, lotions, lipstick, glitter’, |
| ’pollen, vegetation, botany, spores, palynological, palynology, plants’, |
| ’fingerprint, fingermark, latent, enhancement’, |
| ’rna, body fluid, biomarker’, |
| ’bloodstain, blood, pattern, arterial, bpa’, |
| ’hair, hairs’, |
| ’skin,trace dna, touch dna, wearer, scalp, trace, touch, skin, hands, touched’, |
| ’blood, haemoglobin’, |
| ’remains, skeletal, bones, cadavers, teeth, dental, tooth, dentine, dentistry, dentin, tissue, bone’, |
| ’sperm, seminal, semen, postcoital, vaginal’, |
| ’soils, soil, mineralogy, dust, geology, geoscience’, |
| ’condom, lubricant’, |
| ’tape, adhesive, glue’, |
| ’fingernail, nail’, |
| ’saliva, mouth’, |
| ’fibre, fiber, fabric, textile, cotton, garments, nylon, wool, dyed, polyester, shirt, garment, cloth, clothing, clothes, material’, |
| ’cartridge, ammunition, ammunitions, bullet, cartridges’, |
| ’laundry, washing’, |
| ’dna’, |
| ’protein, proteomic’, |
| ’gsr, gunshot residue, gunpowder, elemental, residues, compounds’, |
| ’explosive device, bomb, explosives’, |
| ’arson, ignitable liquid, accelerant, petrol, gasoline, fires,hydrocarbons’, |
| ’firearm, handgun, rifle, pistol, shotgun, gun, weapon’, |
| ’cord, rope’, |
| ’digital forensics, mobile phone, computer, device, file, data’, |
| ’psychology, mental trauma, psychiatry’, |
| ’postmortem,autopsy, injury, wound’ |

---

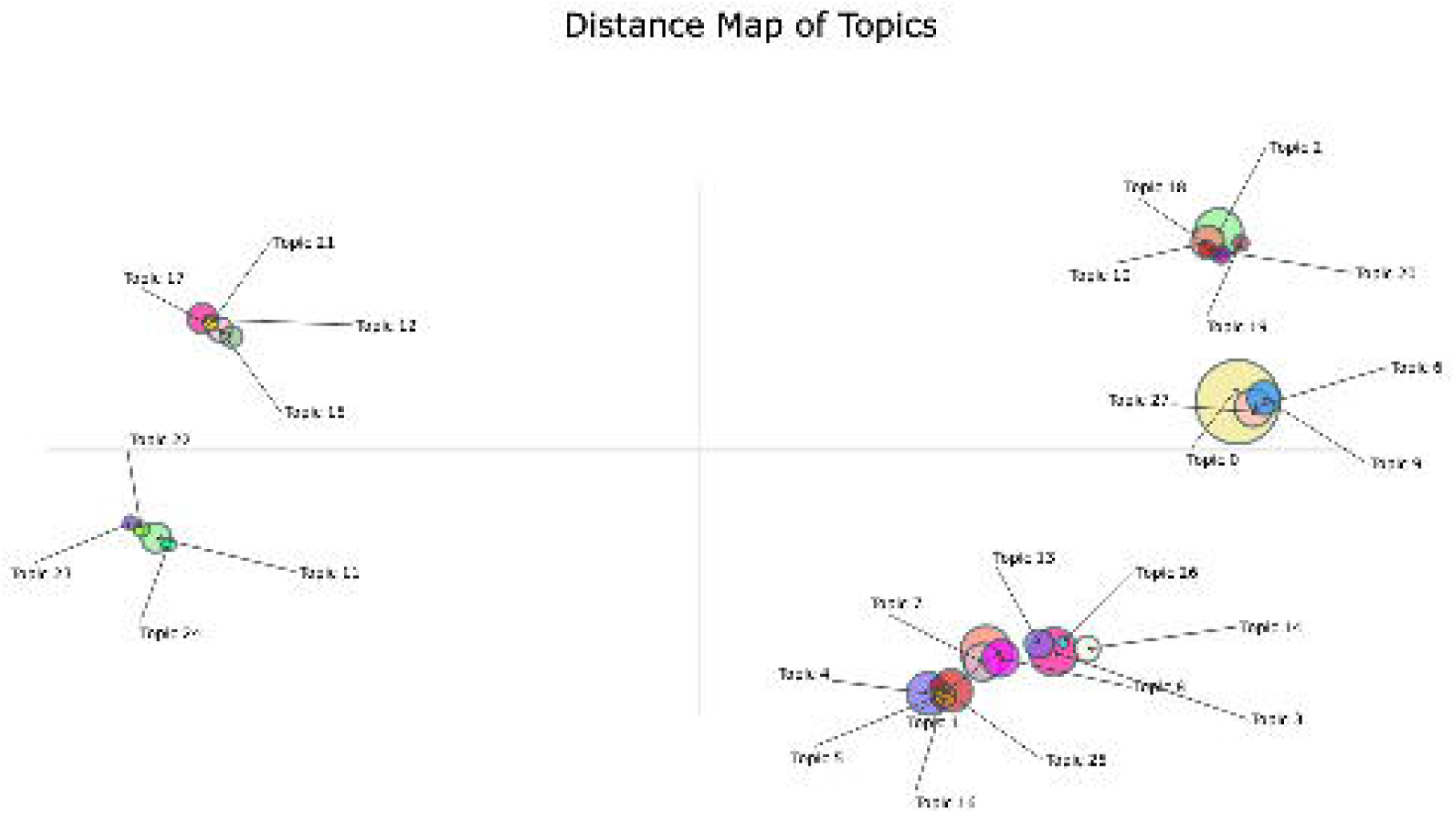

